# Biting rhythm, fecundity and longevity rate of *Aedes albopictus* (Skuse) females from different urbanized settings in Penang Island, Malaysia under uncontrolled laboratory conditions

**DOI:** 10.1101/657023

**Authors:** Nor Atikah Farhah Muhammad, Nur Faeza Abu Kassim, Abdul Hafiz Ab Majid, Hamady Dieng, Silas Wintuma Avicor

## Abstract

Urbanization could potentially modify *Aedes albopictus’* ecology by changing the dynamics of the species, and increasing the abundance of their breeding sites due to environmental changes, and thus contribute to dengue outbreaks. An efficient control of the vector requires a deeper understanding on the biological components of this vector. Thus, this study was conducted to evaluate the biting rhythm, fecundity and longevity rate of *Ae. albopictus* in relation to urbanization strata; urban, suburban and rural areas in Penang Island, Malaysia. The experiments were done in comparison to a laboratory strain. Twenty-four hours biting activity of all the mosquito strains showed a clear bimodal biting activity, with morning and evening twilight peaks. A two-way analysis of variance (ANOVA) found that there was statistically no significant interaction (F(69,192) = 1.337, P > 0.05) between the effects of biting time and mosquito strains. Meanwhile, fecundity rates were shown to be statistically significantly different between mosquito strains (F(3,442) = 10.559, P < 0.05) with urban areas having higher mean number of eggs (mean = 107.69, standard error = 3.98) than suburban (mean = 94.48, standard error = 5.18), and rural areas (mean = 72.52, standard error = 3.87). Longevity rates were significantly higher (F(3,441) = 31.259, P < 0.05) for mosquito strains from urban areas compared to the other strains. These findings would provide crucial and relevant fundamental information to the planning of control program in Malaysia, particularly Penang.

**Author Summary:** Aedes mosquito populations associated with human habitation in urban area do not only have the potential to cause biting nuisance, but also cause significant public health risks through the transmission of dengue virus. The socioeconomic effects of urbanization have been comprehensively studied by socio-ecologists, but the ecological effects and their impact on this vector biology was not known. The authors found that in Penang Island, the mean number of eggs laid per female of *Aedes albopictus* is high in the urban areas than those in suburban and rural areas. The survivorship is high for urban populations parallel to the fecundity rate and apparent biting pattern which is peak at dawn and dusk was noted for all *Ae. albopictus* strains. The changed environment in the urbanized area where more kinds of breeding containers and more blood sources produced by condensed human population supported by warm climate may facilitate larval development, enhance the vector survivorship and its reproductive fitness. These might be the reasons for quick adaptation and susceptibility of *Ae. albopictus* in urban areas. As higher fecundity rate and longer adult survival may enhance disease transmission, this species studied is indeed need high attention in terms of vector control.

## Introduction

Urbanization primarily results in the physical growth of urban areas, leading to environmental changes. It is a global trend that results from economic development. Malaysia is among the fastest growing developing countries in the East [1], and the unprecedented movement of people into these areas is predicted to aggravate in the future [17]. A lot of problems have arisen as a result of urbanization and Malaysia is no exception. Among the problems that often take center stage in urbanization debates include environmental pollution, space, population density and the destruction of natural ecology [20]. The socioeconomic effects of urbanization have been comprehensively studied by socio-ecologists [22, 28, 15]. However, the ecological effects and their impact on vector biology and vector-borne infectious disease transmission remain unclear.

Most of the previous studies were concerned mainly on oviposition ecology from larval habitats and abundance of *Aedes* species in various urbanized settings (13, 20, 29]. However, the biting activity, fecundity and survivorship of individual female *Ae. albopictus* is unknown in response to urbanization level particularly in Penang Island. In order to determine how environmental changes due to urbanization affect the life history traits of *Ae. albopictus* in terms of survival and reproductive fitness, this present study was conducted to determine the biting rhythm, egg production and longevity rate of *Ae. albopictus* strain from Jelutong (urban), Batu Maung (suburban), and Balik Pulau (rural) in Penang Island in comparison to laboratory strain. Since immature mosquitoes are sensitive to environmental changes [18, 24, 20], we hypothesized that urbanization increases *Ae. albopictus* eggs production and survivorship of adult. The findings would provide valuable information on *Ae. albopictus* and would help in improving current vector control and surveillance strategies for dengue that are adapted for specific settings in Penang Island.

## Materials and methods

### Study sites

The ovitrap sampling was conducted in three different areas which represent urban, suburban and rural settings in Penang Island (Fig 1). The distance between each area is approximately 21 km.

**Fig 1.**
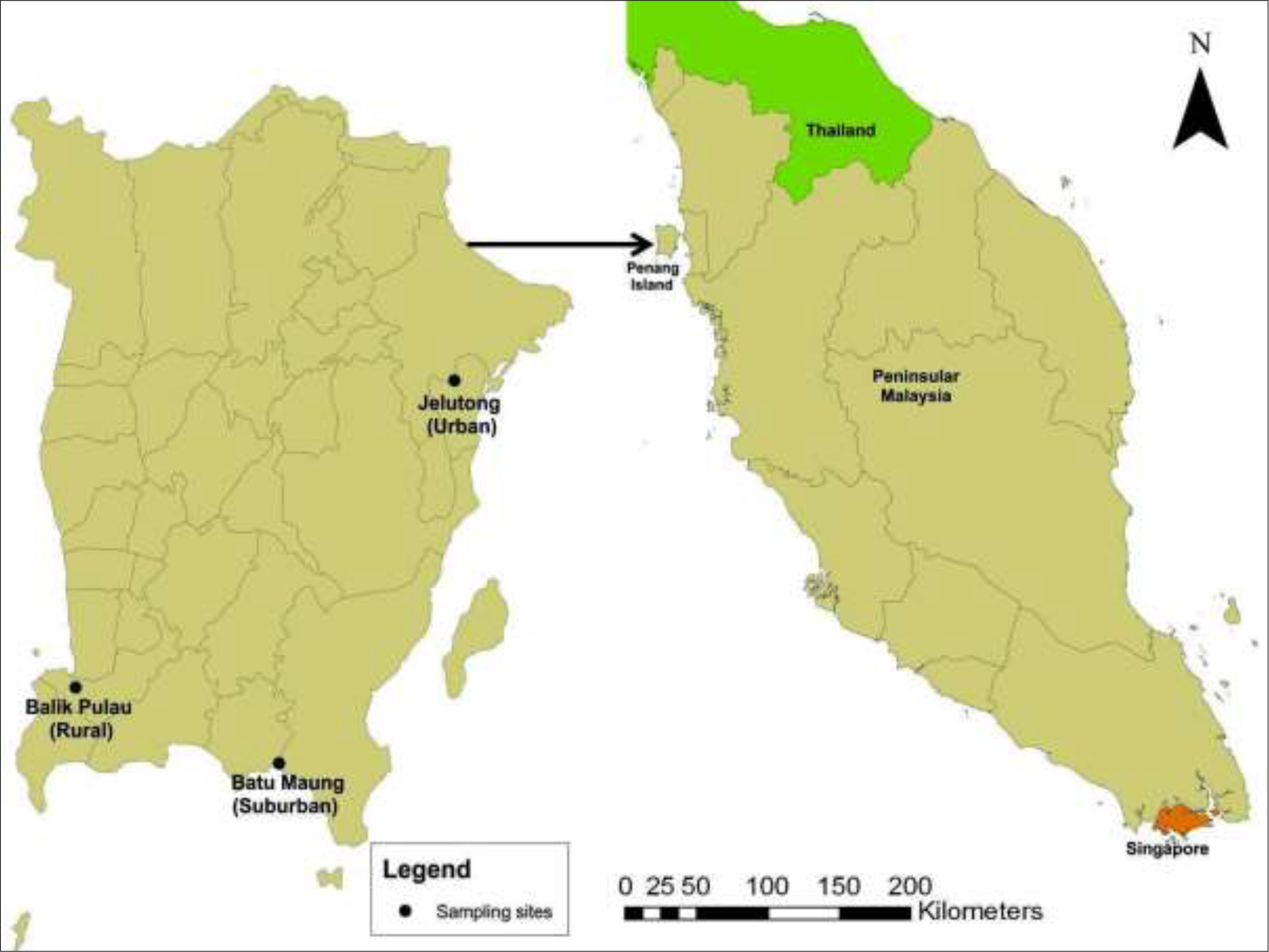
A map of Penang Island showing the location of sites from which *Aedes albopictus* was collected (Source: ArcGIS 10.6 software).

Urban area is an area that is in a city without wooded areas containing a lot of buildings and houses close to each other, high densities of human population, where people often store water for domestic usage due to insufficient water supply (from indoor) (Rahman, 2012). The urban location chosen for this study was Jelutong (5°23’15.9”N 100°18’29.2”E) which is located at the Northeast Penang Island District. It is an urban area which contains little vegetation with the land use types being primarily residential and commercial buildings which are close to each other and public services.

Whereas, suburban refers to a residential area adjacent to the main city area and generally consists of a small number of scattered vegetation, less buildings and population density, but greater house yards in comparison to urban area [26]. The selected suburban location was Batu Maung (5°17’14.5”N 100°16’59.5”E), an area with land use that includes a mixture of residential, manufacturing, and several small fishing villages.

Rural area is considered to be any large and open area, isolated from the city and surrounded with high vegetation cover or small forests. The number of houses is smaller compared to urban and suburban areas due to low human population density. The study site in the rural area was Kampung Pulau Betung in Balik Pulau (5°18’13.4”N 100°11’54.0”E) which is located near a fishing ground, with an economy based on catching fish and harvesting seafood.

Both Batu Maung and Balik Pulau are located at the Southwest Penang Island District. All sample collections was done on public land meanwhile the laboratory strain of *Ae. albopictus* mosquitoes were retrieved from the Vector Control Research Unit (VCRU), USM.

### Mosquito sampling and rearing methods

In this study, *Ae. albopictus* were collected as larvae (fourth instar) and pupae from the three study sites. A total of 30 ovitraps were deployed randomly at two meters apart throughout the respective study areas at shaded sites to maximize the attractiveness of females to oviposit [23]. Each of the ovitraps was filled with approximately 250 ml of chlorine-free water and a paddle (10 cm × 2.5 × 0.3 cm) was placed as an oviposition substrate for the mosquitoes to lay eggs. Five days later, samples from the ovitraps were placed into plastic bottles and brought back to the insectary for rearing. Only larvae (fourth instar) and pupae samples were transferred into paper cups as temporary containers containing water from the ovitraps used and the top of the cup surface covered with nylon netting. The pupae were reared until adult emergence. The eggs from the laboratory strain which were retrieved from the VCRU were reared until adult emergence to be used in the experiment as well. Mosquitoes were reared under 28.6 ± 1.8°C and relative humidity of 65-80%.

Throughout this study, temperature and relative humidity in the insectary were allowed to fluctuate with the weather outside which is an uncontrolled condition. Photoperiod was also unregulated and changed with the surrounding environment. Windows were opened and neither temperature controller nor air conditioner was used during this study.

### Mosquito colonies

Upon emergence, adults were identified based on basic identification keys (dorsal features) following Rattanarithikul and Panthusiri [27]. Only *Aedes albopictus* species were transferred into a standard mosquito rearing cage (30 cm × 30 cm × 30 cm) covered with nylon netting accordingly. For each mosquito strain (urban, suburban, rural and laboratory strain), 40 females and 40 males aged 3 to 5 days old were placed simultaneously in a cage. The mosquitoes were given access to a small cotton wool soaked in 10% sucrose solution on the first day of their emergence and were replaced every two to three days in order to avoid any fungal growth. The mosquitoes were allowed to freely mate for two days, and thereafter starved for a brief period (12 h) [14] before the experiments were executed.

### Biting rhythm experimental design

The first experiment was conducted to determine the biting rhythm of *Aedes albopictus* females from the four different strains within 24 hours. On the day of the experiment, the female mosquitoes were offered a restrained mouse in a narrow and fine wire-mesh cage, at 20:30 h, which is 1 h after sunset when *Ae. albopictus* is reported to be almost inactive [16, 21]. The experiment was ended at 20:30 h on the next day. In approval of animal ethics from The Animal Ethics Committee, USM, the mosquitoes were offered with continuous blood meal for 24 hours, and the mouse was changed at every four hours. Engorgement was checked at 24 time points (21:30, 22:30, 23:30, 24:30, 01:30, 02:30, 03:30, 04:30, 05:30, 06:30, 07.30, 08:30, 09:30, 10:30, 11:30, 12:30, 13:30, 14:30, 15:30, 16:30, 17:30, 18:30, 19:30 and 20:30) and the feeding time was recorded for each female. The fully-engorged females resting on the wall of the cage were removed and transferred into another cage in order to avoid confusion. The experiment was repeated for three times as replicate for each mosquito strain (80 mosquitoes per replicate; 40 females and 40 males).

### Fecundity and longevity experimental design

The second and third experiments were conducted to determine the egg production as well as the longevity rate of *Aedes albopictus* females from the four different strains. The female mosquitoes as mentioned in the **sub-section Mosquito colonies** were offered a restrained mouse in a narrow and fine wire-mesh cage for one hour; 18:00-19:00 hour, the closest time to the peak biting activity of *Ae. albopictus* in the field [3].

A day after, females were considered gravid [23]. Every gravid female was then placed singly into an oviposition cage (1.3 L) consisting of modified plastic bottle supplied with a small cotton wool soaked in 10% sucrose solution through a thin nylon cloth covering the top end of the bottle (Fig 2). Cotton wools were replaced every two to three days. A small disposable plastic cup filled with 30 ml of chlorine-free water and lined with filter paper as an oviposition substrate was placed in each cage to provide females with sites for egg deposition. The filter paper was folded into a double-chambered cone and placed so that the mosquitoes could lay their eggs inside or outside the cone [14]. The separation of the females into individual oviposition cages facilitated the recording of the number of eggs produced by each individual female.

**Fig 2.**
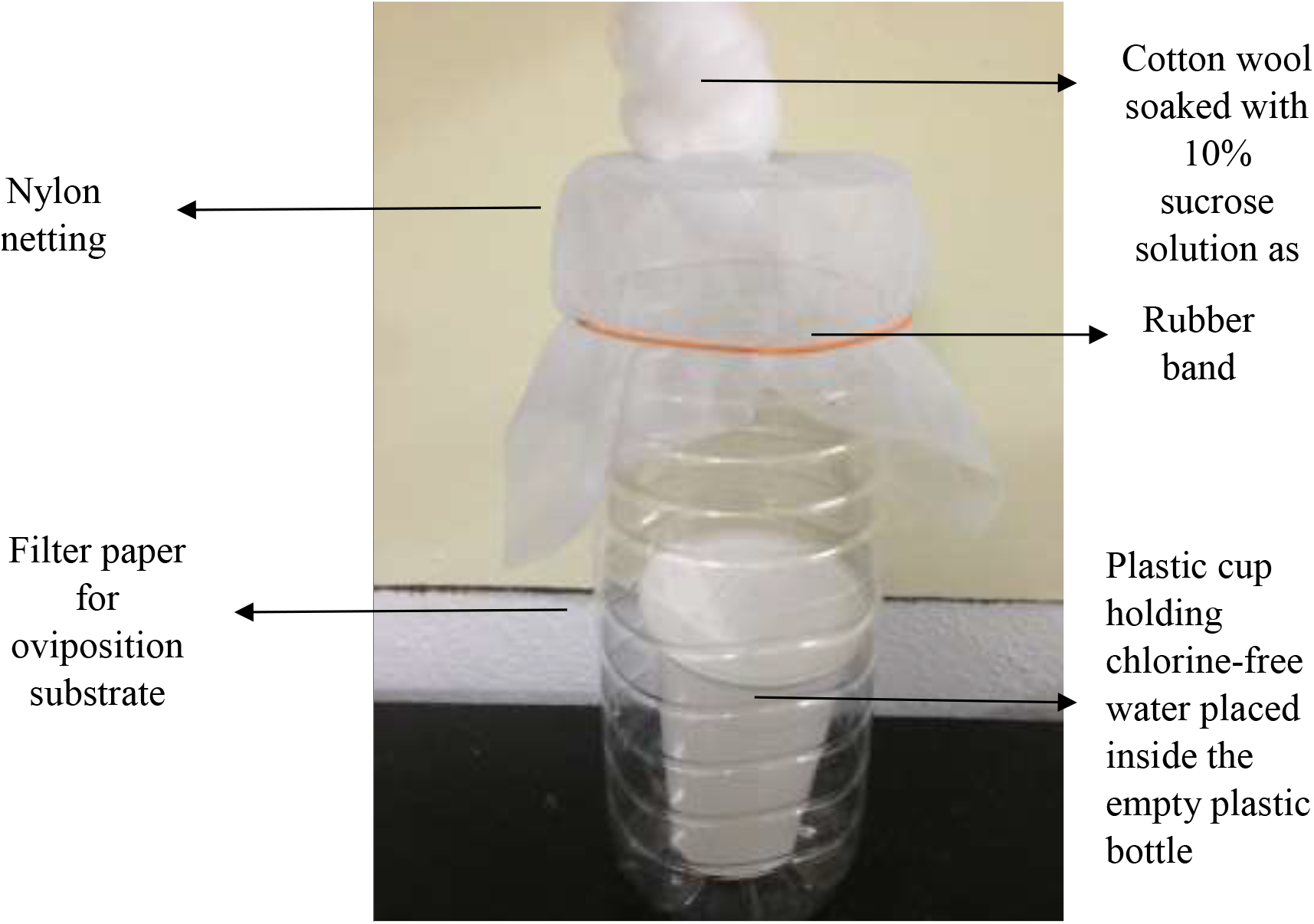
Modified oviposition cage.

Two days later, egg deposition was checked daily and, filter paper with eggs was removed. Eggs were counted under dissecting microscope, and a new filter paper was placed in each cage. This process was repeated until their death. In order to determine the longevity rate of the *Ae. albopictus* females, the dead mosquitoes were recorded and removed from the cage daily. The experiment was repeated three times as replicate for each mosquito strains (80 mosquitoes per replicate; 40 females and 40 males).

Throughout the series of experiments involved in this study, temperature and relative humidity in the insectary were allowed to fluctuate with the weather outside which were uncontrolled conditions in order to imitate the actual surrounding environment of mosquitoes [23]. Photoperiod was also unregulated and changed with the surrounding environment. Windows were opened and neither temperature controller nor air conditioner was used during this study.

### Data analysis

The analyses were conducted using Statistical Package for the Social Sciences (SPSS) version 20.0. Data were tested for normality using Shapiro-Wilk statistic (P > 0.05). When data was confirmed for normality it was further analyzed but if the data were not normally distributed, data transformation is needed to fulfill the assumption of required tests.

For biting rhythm, the number of fully-engorged mosquitoes (dependent variable) from four different strains in relation to blood feeding time (independent variables) were subjected to two-way analysis of variance (ANOVA). A two-way ANOVA was conducted to examine the effect of time and mosquito strain on biting rhythms of *Ae. albopictus* females.

Whereas, for fecundity rate, the number of eggs produced per mosquito (dependent variable) was used to compare between mosquito strains (independent variable) and were analyzed using one-way ANOVA. A one-way ANOVA was tested to determine if there were any differences in the fecundity rates for the four different mosquito strains. Tukey’s multiple comparison test was further analyzed to determine significant differences between strains.

Meanwhile, Kaplan-Meier survival analysis along with a log-rank test was performed to estimate the longevity rate of *Ae. albopictus* females from each strain. A log-rank test was run to determine if there were differences in the survival distribution for the four different mosquito strains.

## Results

### Biting rhythm of Aedes albopictus females from four different strains; urban, suburban, rural, and laboratory

Figure 3 shows the result of the 24 h biting cycle experiment performed under uncontrolled laboratory conditions. The experiment revealed two activity peaks for *Aedes albopictus*, coinciding with crepuscular dawn, 06:30-09:30 hours, and dusk, 18:30-20:30 hours for all four mosquito strains.

**Fig 3.**
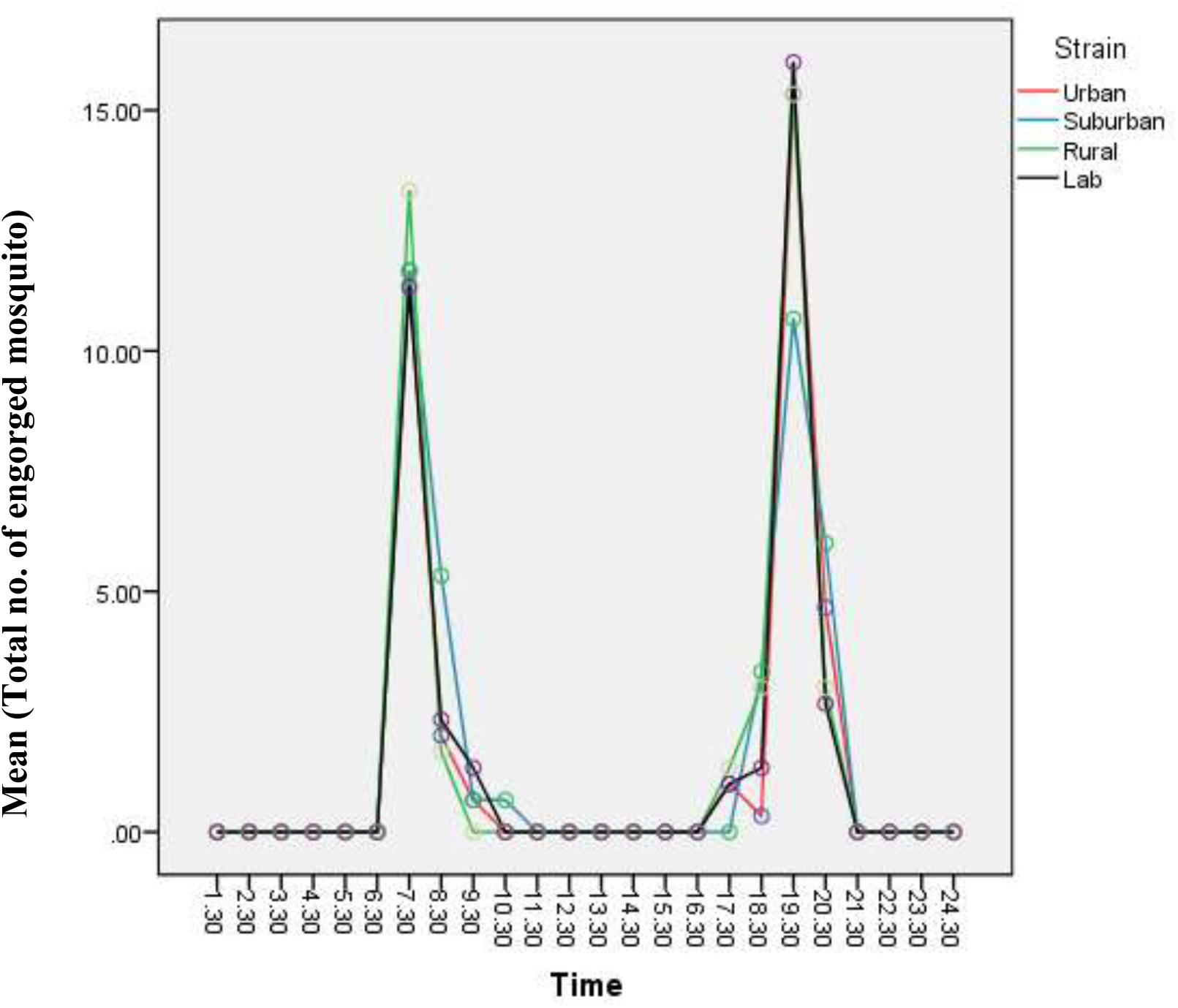
Biting rhythm of *Aedes albopictus* females from four strains; urban, suburban, rural, and laboratory.

There was statistically no significant interaction between the effects of biting time and mosquito strains (F(69,192) = 1.337, P > 0.05) on number of engorged *Ae. albopictus* female mosquitoes. Thus, *Ae. albopictus* biting cycle remained as a bimodal pattern regardless of the strain of mosquito [16, 2]. Aside from bimodal, the term is most often described as crepuscular in which the insects are those that are active primarily during twilight [25].

### Fecundity rate of Aedes albopictus females from four different strains; urban, suburban, rural, and laboratory

Based on the fecundity test conducted, the mean number of eggs produced by *Ae. albopictus* increased from laboratory to urban areas. Referring to Fig 4, females from the urban area showed the highest mean recorded with 107.69 ± 3.98 number of eggs produced per mosquito than those in suburban (94.48 ± 5.18), rural (72.52 ± 3.87) and laboratory strain (53.65 ± 2.34).

**Fig 4.**
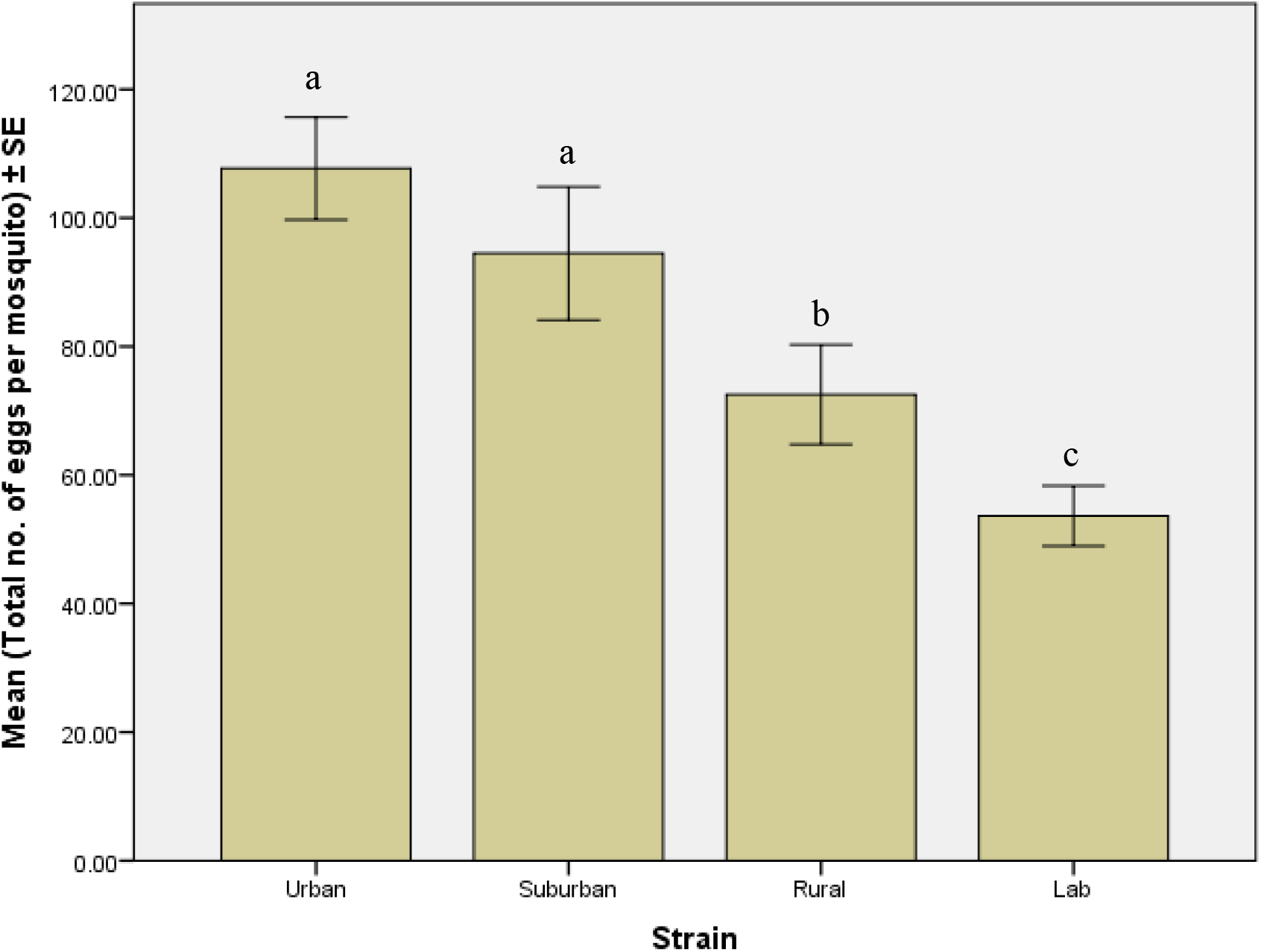
Total number of eggs produced by individual *Aedes albopictus* female in three different strains. **the same letter indicated no significant difference between strain (P > 0.05) **the different letter indicated significant difference between strain (P < 0.05)

There was a significant difference in egg production between strains (F(3,442) = 10.5, P = < 0.05). Tukey’s multiple comparison test further showed that the fecundity rates for urban and suburban mosquito strains were significantly different from rural and laboratory strains (P < 0.05).

### Longevity rate of Aedes albopictus females from four different strains; urban, suburban, rural and laboratory

The survival analysis shows distinct differences of the longevity rate of *Ae. albopictus* females between four strains (urban, suburban, rural and laboratory strain). Referring to Fig 5, females from the urban area showed the highest mean days of lifespan with 25.29 ± 1.10 than those in suburban (23.12 ± 0.97), in rural (18.60 ± 1.14), and laboratory strain (11.89 ± 0.90).

**Fig 5.**
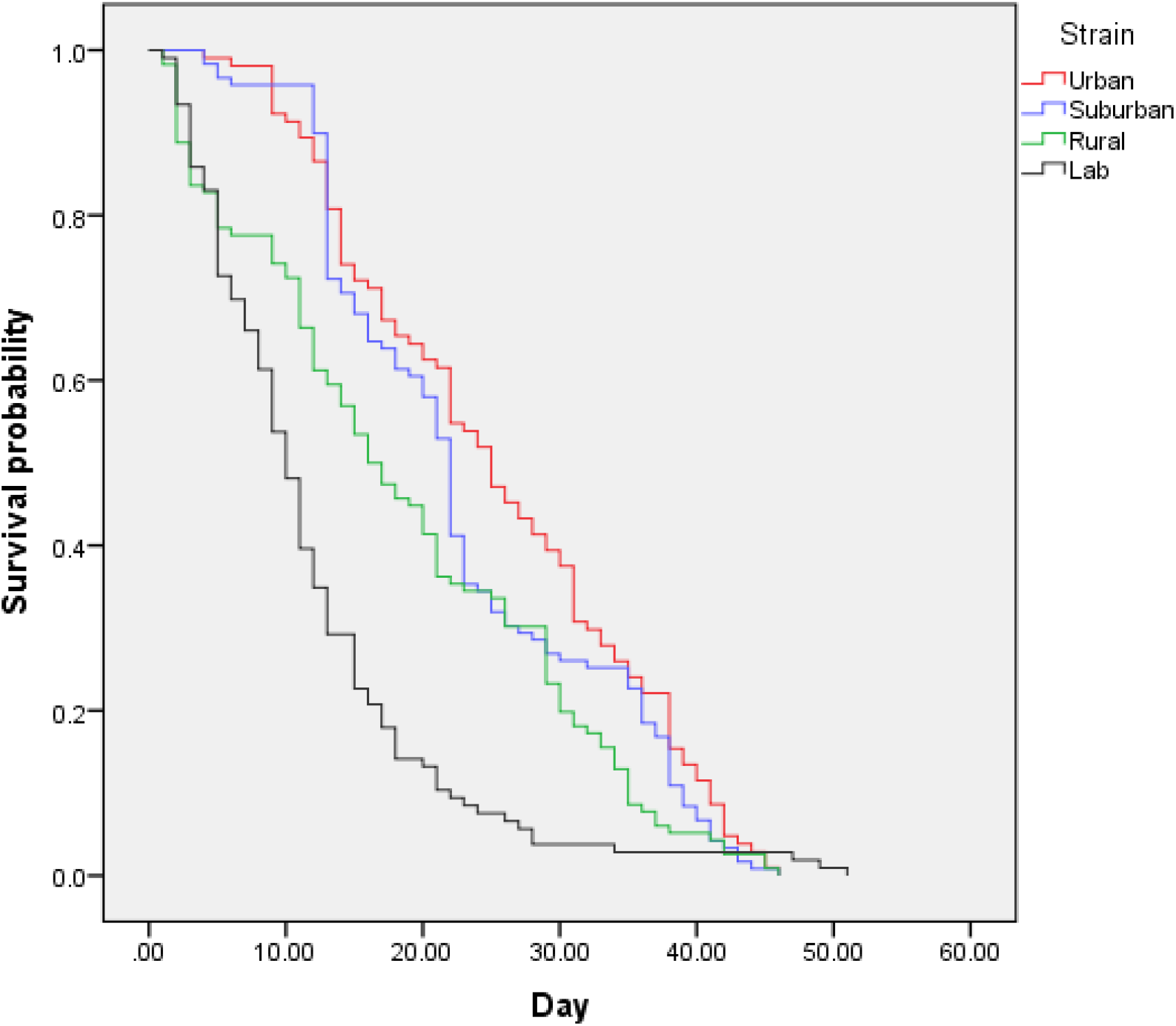
Cumulative survival probability of *Aedes albopictus* females from four different strains.

A log-rank test showed significant difference in the survival distributions for the four different mosquito strains (*X*^*2*^ = 72.28, d.f. = 3, P < 0.05).

## Discussion

Based on the 24 h biting activity of all mosquito strains, *Ae. albopictus* females showed a clear bimodal activity, with morning and evening twilight peaks, as observed elsewhere in Southeast Asia through human landing catch technique in the field [16,2] However, we observed the biting activity under uncontrolled laboratory conditions, where *Ae. albopictus* females were offered with restrained mouse as their blood meal at 20:30 h. When we checked for engorgement at the next hour in accordance to the time points, surprisingly, the blood-feeding activity as well as host-seeking activity was observed to be inactive. The same situation happened for almost ten hours of the experiment (21:30 h to 06:30 h). No engorged mosquitoes resting on the cage wall were recorded within that duration.

This behavior might simply explain the biting nature of this species. According to [19], the nocturnal host-seeking activity positively correlated with the increasing light intensity as *Ae. albopictus* is sensitive to dim light. In their study, they found out that the host-seeking activity for this species is deactivated in complete darkness even during the daytime, irrespective of their increasing flight activity controlled by their intrinsic circadian rhythms [19]. Their host-seeking activity changes when the light intensity during the scotophase was changed daily from 0 to 100 lux [19]. The rapid drop in activity level following the onset of darkness suggested that light promotes activity or, less likely, that the absence of light reduces the activity. Light may have a direct effect in determining the amount of activity and indirect effect through setting the phase of the endogenous rhythm [30]. The nocturnal feeding might not be due to the evolutional adaptation to light, but by the intrinsic reaction of mosquito to light [19]. Thus, the potential risks of transmission possibilities of dengue virus exist wherever the vector inhabits (urban, suburban or rural areas).

The previous studies were concerned mainly on oviposition ecology and abundance of species by urbanization [13, 20, 29] without knowing how many eggs one female could produce in a lifetime. By increasing eggs production by *Aedes* mosquitoes, urbanization could potentially worsen the epidemic risk factors for arboviruses. An increase in *Ae. albopictus* species prevalence and abundance by urbanization was reported by [20] where this phenomenon is probably due to elevated numbers of *Aedes* breeding sites such as tires, discarded cans or water storage containers, provided by urbanizing environment [20]. They reported that an urbanized environment accelerates *Aedes* mosquito development and survivorship. A deeper understanding of the modifications induced by urbanization in the ecology of *Ae. albopictus* is indispensable in order to improvise the vector control strategies in the future.

In the present study, fecundity or mean number of eggs laid per female of *Ae. albopictus* in the first gonotrophic cycle was high in the urban areas (107.69 ± 3.98) than those in suburban (94.48 ± 5.18), and rural areas (72.52 ± 3.87). This finding was supported by a study done by [12]. In their study, they found that *Aedes aegypti* which exist in forests had lower reproductive potentials than the typical form which lives in urban and suburban areas. Therefore, they speculated that the forest probably provides a more homogenous environmental stress than the urban habitat, where urban populations have adapted to a fluctuating environment which is probably related to variation in the availability of larval habitats, food source, temperature as well as humidity range. Egg production in the field is dependent on environmental factors such as temperature and relative humidity, since these factors influence the survival of adult mosquitoes. Warmer temperature will accelerate the digestion of blood meals taken by mosquitoes, leading to increased human biting frequency and increased fecundity rates and reproductive fitness [5]. Apart from that, the number of eggs laid by *Ae. albopictus* female also depends on the physiological age, the body weight after emergence and particularly the size of the blood meal [9]. The development of the immature stages may influence egg production too. An optimum condition during the growth of immature will result in larger and healthier adults who can consume more blood from the hosts [23]. The amount of blood imbibed and the blood digestion rate by the female determines the number of eggs produced. More eggs will be produced when an adequate amount of blood is consumed and rapidly digested [7].

These results reflected in the abundance of pupal and larval density in urban area rather than suburban and rural areas [6, 20]. This might be due to the fact that urban areas had less predators, more nutrition from a “dirtier” environment, or even less drift from agricultural insecticides [20]. Pupal productivity is a good indicator of the abundance of adult mosquitoes [8, 10, 11].

Higher mosquito fecundity does not necessarily lead to increased disease transmission if adult mosquitoes have a very short life span. This is because longevity must be sufficiently long in order to allow for the development of pathogens in their bodies to be completed. We found that female mosquitoes in urban areas had the longest life span compared to the other two areas. This result may be due to environmental factors such as temperature as well as humidity, where the average temperature in urban areas is higher than in suburban and rural areas [20]. This findings was similar with the previous study done by [4], where they found that the median survival of *Anopheles arabiensis* in the deforested area was 49 to 55% higher than those in the forested area and the net reproductive rate of female mosquitoes in the deforested area was 1.7 to 2.6 fold higher than that in the forested area in the East of Africa. Therefore, they concluded that deforestation or urbanization enhanced the survivorship of adult mosquitoes [4].

Longer adult longevity may enhance disease transmission, although the exact correlation between vector capacity and adult life span needs to be further explored. In this study, we fed the female mosquitoes with 10% sucrose solution without blood, which might have led to exerted stress on the females during gonotrophic cycle and affected the longevity of the female mosquitoes [23]. Thus, we predicted that the females will live much longer if they were given access to blood like in the natural environment. Overall, the findings clearly showed how *Aedes* mosquitoes are better adapted to urban environment.

The fecundity and longevity rate of laboratory strain mosquitoes was recorded as the lowest in comparison to the other wild strains. This might be due to the environmental stress as the experiment was conducted under uncontrolled laboratory condition, where they are not used to.

## Conclusion

In conclusion, the results of this study indicated that urbanization might have a significant impact on the ecology as well as biology of *Aedes albopictus*. In the urbanizing and urbanized area, the changed environment probably serves as a suitable place for the growth and development of *Ae. albopictus;* the condensed human population produced more kinds of containers for larval habitats and more blood sources for adult reproduction. Warm climate may facilitate larval development, enhance the vector survivorship and reproductive fitness. These might be the reasons for quick adaptation and susceptibility of *Ae. albopictus* in urban areas. Urbanization in Penang Island has exhibited strong positive effects on the reproductive and survivorship of adult mosquitoes through effects on the microclimatic condition of the mosquitoes, hence encourage dengue outbreak. The implications of these findings are that, if the current trends of urbanization continue in the rural area, *Ae. albopictus* could adapt to the changing environment and proliferate in the rural areas, resulting in vast dengue outbreak in the future. These findings would provide crucial information to the dengue vector control program and may be helpful to vector control experts in predicting outbreaks which could worsen if urbanization occurs without limits.

## Acknowledgements

The authors are grateful to the Vector Control Research Unit, Universiti Sains Malaysia and the respective communities for logistical support and sample collection respectively. Thanks also to Dr. Azimah Abd Rahman for her kind help. This study was funded by the Short Term Grant (304/PBIOLOGI/6313064) awarded by the Division of Research and Innovation, Universiti Sains Malaysia and External Agency Grant of Universiti Sains Malaysia, the Ministry of Natural Resources and Environment – NRE (304/PBIOLOGI/650880/K130).

